# An Implementation of Empirical Bayesian Inference and Non-Null Bootstrapping for Threshold Selection and Power Estimation in Multiple and Single Statistical Testing

**DOI:** 10.1101/342964

**Authors:** Bahman Nasseroleslami

## Abstract

The majority of conclusions and interpretations in quantitative sciences such as neuroscience are based on statistical tests. However, the statistical inferences, especially in multivariate analyses, commonly rely on the p-values, but not on more expressive measures such as posterior probabilities, false discovery rates (FDR) and statistical power (1 − *β*). The aim of this report is to make these statistical measures further accessible in single and multiple statistical testing. For multiple testing, the Empirical Bayesian Inference (Efron et al., 2001; Efron, 2007) was implemented using non-parametric test statistics (e.g. the Area Under the Curve of the Receiving Operator Characteristics Curve or Spearman’s rank correlation) and Gaussian Mixture Model estimation of the probability density function of the original and bootstrapped data. For single statistical tests, the same test statistics were used to construct and estimate the null and non-null probability density functions using bootstrapping under null and non-null grouping assumptions. Simulations were used to test the reliability of the results under a wide range of conditions. The results show conformity to the real truth in the simulated conditions, which is held under various conditions imposed on the simulated data. The open-source MATLAB codes are provided and the utility of the approach has been exemplified and discussed for real-world electroencephalographic signals. This implementation of Empirical Bayesian Inference and informed selection of statistical thresholds are expected to facilitate more realistic scientific deductions in versatile fields, especially in neuroscience, neural signal analysis and neuro-imaging.

## 1 Introduction

The majority, if not all, of the conclusions and interpretations in quantitative sciences, especially in neuroscience and neuro-imaging, are based on statistical tests. While the traditional hypothesis tests based on p-values are still dominant, there has been legitimate remarks on the need for more reliable and thorough statistical procedures and practices (Nuzzo, 2014). For statistical inference, it is therefore vital to make accessible more meaningful statistical measures, including Bayesian posterior probabilities, false discovery rates (FDR), and statistical power (1 − *β*). These measures are especially useful in statistical inferences involving high-dimensional neuroimaging or neural signal connectivity data.

The traditional p-value represents the probability of the observed data under the null hypothesis. The lower the p-value, the more unlikely the null hypothesis is. Therefore, indirectly, p-values lower than the threshold value *α* (commonly taken as 0.05 by convention) are used to reject the null hypothesis and support the alternative hypothesis; however, this view has been subject to criticism (Amrhein, Greenland, and McShane, 2019; Nuzzo, 2014). The low p-values indicate the low probability of false positives (Type I statistical error). Therefore, a more reliable measure for statistical inference of the observed effects is probably the probability of true positives, which corresponds to the statistical power (a probability closely related to the reproducibility). Statitical power is 1−*β* where *β* is the Type II statistical error, false negative. This basic concepts can be reviewed from texts in statistics for classic univariate statistics (Moore, McCabe, and Craig, 2010). The statistical power would provide further insight in situations where the Bayesian Posterior probability (probability of the effect given the observed data) cannot be calculated - the most common reason being the unknown value for the prior probability (see Methods). In more practical applications, e.g. multidimensional analysis of neural signals, multivariate statistics are needed for an informed statistical inference. In such situations, measures such as False Discovery Rate (FDR; Benjamini and Hochberg, 1995; Benjamini, Krieger, and Yekutieli, 2006), that refers to the proportion of the detections that are falsely identified; as well as multivariate power (the proportion of the actually affected variables that were successfully detected by the test), are more instrumental for scientific inference. Importantly, the presence of several variables provides opportunity for inferring the prior probability from the data, and hence, estimating the posterior probability (Efron, Tibshirani, Storey, and Tusher, 2001).

Empirical Bayesian Inference (EBI) has shown promise in large-scale between-group comparisons (Efron, 2004, 2007b), especially in genomics (Efron et al., 2001) and to some extent in the applications of neuroelectric signal and connectivity analysis (Singh, Asoh, Takeda, and Phillips, 2015). In EBI, constant prior probabilities are estimated from the data in large-scale multi-variable inferences or hypothesis testing and these priors are subsequently used to find the posterior probabilities using the estimated probability density functions of the pooled test statistics and the null distribution. It is possible to relate the posterior probabilities to frequentist concepts such as FDR, as well as power. While the theoretical framework is adequately established, the existing mathematical and numerical implementations (Efron, 2007b) are only suitable for specialised applications (i.e. statistical genetics, where only a small fraction of the tests are real findings), and some of the essential measures such as FDR and power are not immediately available for informed threshold selection. ***Consequently, there is no implementation of EBI that works for any data in unknown arbitrary conditions.*** From a practical viewpoint, the existing software package *locfdr* in R (R Core Team, 2016) may require selection of several parameters and is not immediately available for neuro-electro-magnetic signals (e.g. EEG and EMG) and connectivity analysis in packages such as FieldTrip (Oostenveld, Fries, Maris, & Schoffelen, 2010) or for neuroimaging analysis in packages such as SPM (Ashburner, 2012; Friston et al., 1994). The commonly applied methods such as cluster-based permutation (Bullmore et al., 1999; Maris, Schoffelen, and Fries, 2007), do control for FDR at specific levels; however, they suffer from 4 main constraints: 1) They rely on the implicit assumption that the significant detections are spatially clustered and therefore playing down the smaller spatial clusters, 2) the lack of a criterion to define clusters of variables, leading to arbitrary cluster definitions, 3) inability to estimate the posterior probability or statistical power, and 4) difficulty in evaluation and deciding on a significance thresholds. Consequently, there is the need for new implementations to facilitate the application of EBI that work in wider range of situations (e.g. small or large proportions of test variables belonging to the affected group), to more explicitly relate the posterior probabilities to FDR and power (allowing informed decision on threshold selection), and to intrinsically account for data with non-normal distributions.

Such informed selection of statistical threshold is challenging also in complex statistical inferences (e.g. with non-normal data distributions) involving single or only a few comparisons or inferences. It would be desirable to similarly select a threshold value for the test statistic that corresponds to a known combination of Type I (*α*) and Type II (*β*) errors in a single comparison.

Here, these needs are address using an implementation of EBI using non-parametric test statistics, Gaussian Mixture Models and null bootstrapping. This implementation readily handles one-sample, two-sample (between-group comparison) and correlation problems in multi-dimensional data with arbitrary distributions, which is usable for a wide range of applications. Furthermore, for threshold selection in univariate testing (in the absence of prior probabilities), the non-null distribution is estimated using a non-null bootstrapping. This approach approximates the non-null probability density functions in order to enable the threshold selection for a desired combination of *α* and *β* values, regardless of the distribution of data.

In this report, after setting out the mathematical underpinnings of the EBI, the components of the new implementation are explained. These include the selection of test statistic, the estimation of density functions by Gaussian Mixture Models (GMM), and bootstrapping. The implementation is tested against real truth in simulated data in several conditions, and finally demonstrated using experimental neuro-electric data (EEG).

## 2 Methods

### 2.1 Empirical Bayesian Inference (EBI) for Multiple Inferences

#### 2.1.1 EBI framework

EBI, initially used in genomic applications (Efron et al., 2001) was subsequently expanded theoretically and in terms of computational implementation (Efron, 2004, 2007a). Here, the fundamentals are briefly explained. Throughout the report, the procedures are explained for mass univariate two-sample comparison problems as an exemplary scenario. However, the procedures are equally applicable to other problems, e.g. one-sample / paired problems or correlation analysis (discussed in the Appendices).

Suppose *X*_*ij*_, where *i* = 1…*m, j* = 1*…N* represents *N* variables or *N*-dimensional data sampled from *m* observations/subjects. In the case of two-sample data, *X*_*ij*_ represents pooled data. The grouping information of data is represented by *g*_*i*_, which, for a two-sample (2-group) comparison is a binary choice (0 or 1). Statistical testing, performed independently in each variables according to the grouping information (e.g. between-group comparisons), using the test statistic *z*_*j*_, yields *N* values. The probability density function of *z*_*j*_, i.e. the probability of data given the hypotheses, is denoted by:

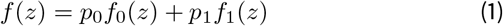

where *p*_0_ and *p*_1_ are the prior probabilities of the null and non-null hypotheses (*p*_1_ = 1 − *p*_0_) and *f*_0_(*z*) and *f*_1_(*z*) are the probability density functions of *z* unwder the null and non-null (grouping) assumptions, respectively. The posterior probability, i.e. the probability of the hypotheses given the data, are subsequently given by:

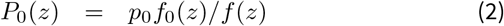

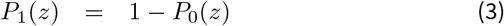

Comparison of the posterior probability of the non-null hypothesis *P*_1_(*z*_*j*_) against a threshold *P*_*crit*_ provides a Bayesian inference, as well as subsequent frequentist quantities such as local false discovery rate, *fdr*_*loc*_(*z*) = *P*_0_(*z*), Type I error *α*, Type II error *β*, and the FDR value pertaining to the chosen *P_crit_*. The original formulation of EBI includes several stages for estimating the posterior probabilities: First, applying a measure of between-group difference (e.g. Student’s *t*-statistic or *p*-values) and transforming the values to normal (e.g. by inverse normal cumulative distribution function) to build *z*_*j*_ values; second, estimation of *f* (*z*) from the *z*_*j*_ histogram; third, estimating *f*_0_(*z*) by theoretical assumptions on the distribution of *z*_*j*_ or bootstrapping; forth, estimation of null prior *p*_0_, usually through the assumption that *f*_1_(*argmax*_*z*_*f*_0_(*z*)) = 0; and finally, *P*_1_(*z*) is found by equations (2) and (3). Here, we will explain the details of each stage for the new implementation.

#### 2.1.2 Test Statistic

Instead of using Student’s *t*-statistic or a *p*-value which reflects the difference of the means of two groups, here a non-parametric measure was used as a test statistic. The Area Under the Receiver Operating Curve (AUROC), *A*, is closely related to the Mann-Whitney *U* statistic. AUROC is the probability of data in one group being larger (or smaller) than the other group (*Pr*(*X*_*g*=0_) < *Pr*(*X*_*g*=1_)); hence it is considerably independent of the distribution of the original data, as well as any measure of centrality (e.g. mean or median) for comparison in parametric testing (e.g the comparison of means as statistic in using t-tests). This has been thoroughly discussed elsewhere (Zhou, McClish, and Obuchowski, 2009). AUROC was therefore taken as the test statistic for comparing the *m* data points in the two groups for each comparison of the *N* variables.

It is noteworthy that while AUROC is independent of the underlying distribution of the data, the data in all *N* variables should come from the same null and alternative (non-null) distributions. The AUROC distributions depend on the number of data in the first and second group, as well as the distribution of the original data. Therefore, the number of data points (e.g. individual subjects in a group comparison) in each group should be (ideally) the same for all variables, and all of them should come from the same arbitrary distribution (e.g. normal, Beta, Gamma, or uniform distribution). This is especially relevant as the curve fitting for null and mixed density functions, as well as bootstrapping (for construction of null data) rely on pooling data from all variables.

While not essential to transform the AUROC values *A*_*j*_ to normal, it’s beneficial to do so, from a computational perspective. The transformation to normal allows the use of more robust estimation methods such as Gaussian kernel methods that work best in unbounded domain, rather than in bounded ([0 1]) domains. As the AUROC distribution is bounded between 0 and 1, with the expected value of 0.5 under null, a mapping of *v* = 2(*A*_*j*_) − 1 combined with Fisher’s Z-transform, *z*_*i*_ = *arctanh*(*h*) = 0.5*log*_*e*_((*v*) + 1) − 0.5*log*_*e*_(1 − (*v*)) (Fisher, 1915; Zhou et al., 2009), can approximately map the data to normal (Qin and Hotilovac, 2007; Zhou et al., 2009):

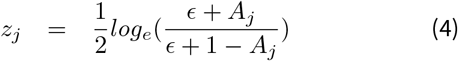

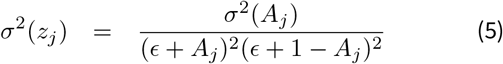

where *σ*^2^ indicates the variance. The tuning parameter *∊* which is added to the classic definition here, serves to limit the extreme *z* values thta would typically span the range (−∞, +∞); hence, facilitating numerical integration in later steps. To limit the *z* values to [−10, 10], *∊* = 2.061 10^−9^ was adopted here. In addition to this, in order to avoid sharp distributions where *AUC* = 1 and *z* = 10 (*AUC* = −1 and *z* = −10), the values larger than 9 (smaller than −9) were redistributed to a truncated normal distribution. The redistribution of *h* extreme values larger than 9, assigned the *i*^*th*^ value to 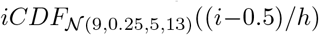, where *iCDF* is the inverse cumulative distribution function and 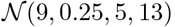, is the normal distribution with mean 9, standard deviation 0.25, truncated between [5, 13]. The values smaller than −9 were similarly reassigned.

#### 2.1.3 Estimating the *f* (*z*) Histogram

Gaussian Mixture Model (GMM) distributions (McLachlan and Peel, 2004) were used to estimate the probability density *f* (*z*), using the pool of *z*_*j*_, *j* = 1…*N*. Using maximum likelihood estimates of GMM parameters, models with increasing number of Gaussian kernels were set for fitting *z*_*j*_ values. The model with minimum Akaike Information Criterion, AIK, (Akaike, 1974) was eventually considered as the preferred fit. This sequential exploration procedure was concluded when the increasing number of kernels yielded 3 consecutive increases in the AIK. A similar approach has been previously used (Le, Pan, and Lin, 2003) in statistical genetics applications, but not in the context of EBI.

#### 2.1.4 Estimating the Null Distribution *f*_0_(*z*)

For robust estimation of the null distribution, the data labels *g*_*i*_ were set for *B*_0_ times re-sampling with substitution; for each set of the obtained resampled labels, the procedures for the original data and labels were applied to yield *A*_*j*_ and subsequently the *z_j_* values. The data from all the *B*_0_ bootstraps and all the *N* tested variables were used for pooling to estimate the null distribution. For computational efficiency, it is helpful to sparse the null distribution. In this implementation, when the null data points exceeded 20000, the null data were sorted and only the every *sk* data values were kept for GMM estimation (*sk*: the integer multiples of 20000 in the number of null data values). Using similar GMM estimation as for *f* (*z*), the null distribution *f*_0_(*z*) was estimated.

#### 2.1.5 Estimating the prior *p*_0_

The approach used by EBI for estimation of the prior *p*_0_, relieson the key assumption that at maximum (peak) value of *f*_0_(*z*), the value of *f*_1_(*z*) is zero. Due to the smooth and reliable estimation of *f*_0_(*z*) and *f* (*z*) by AIK-guided GMM fits, it is possbile to directly use the values of the estimated probability density functions to find the prior *p*_0_:

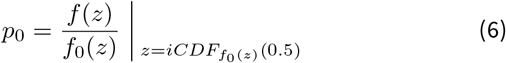

where 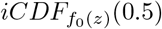 is the *z* value at which the Cumulative Density Function (CDF) of *f*_0_(*z*) is 0.5, which is the inverse CDF of 0.5, i.e. the median of the null data.

#### 2.1.6 Estimating the Posterior *P*_0_(*z*)

Given the estimates of *p*_0_, *f*_0_(*z*) and *f* (*z*), the calculation of posterior probabilities from equations (2) and (3) are straightforward. A bound between 0 and 1 was considered to protect against numerical instability at very small probability values.

#### 2.1.7 Estimating FDR and Power (1 − *β*)

Following the calculation of *p*_0_, *p*_1_, *f*_0_(*z*), *f*_1_(*z*), *f* (*z*), *P*_0_(*z*) and *P*_1_(*z*), the Type I error *α*, Type II error *β*, power (1−*β*) and FDR (*q*) can be found by numerical integrations. This is achieved by using a decision threshold value (see Section 2.1.8 for how this threshold is decided on) on either of these measures to infer which variables do or do not show an interesting effect. For a given decision threshold on *α*, *β*, or *q*, and a corresponding criterion on the Posterior, *P*_*cr*_, we may write:

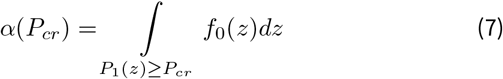

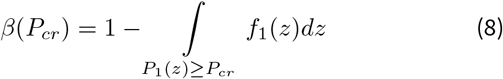

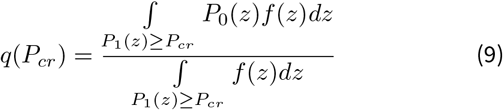

as the parameter *∊* in section 2.1.2 limits the values of *z*, the integration would suffice to take place in the range [−20, 20]. Additionally, the global values *α*_*g*_ and *β*_*g*_ show the sperability of the probability density distributions regardless of a chosen *P*_*cr*_:

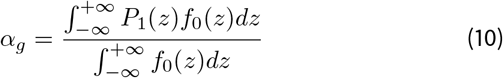

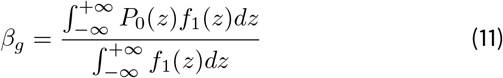

#### 2.1.8 Threshold Selection

The threshold selection and subsequent inference is driven by setting a criterion *P*_*cr*_ on Posterior and comparing the values of *P*_1_(*z*) against this specific *P*_*cr*_, as required in the specific context of application. Alternatively, with the availability of the computed *α*, *β*, *q*, it is possible to find the number of values that are higher than the threshold #{*P*_1_(*z_j_*) ≥ *P*_*cr*_}, and the detection ratio #{*P*_1_(*z*_*j*_) ≥ *P*_*cr*_}/*N* as a function of *P*_1_(*z*); which in turn allows setting the *P*_*cr*_ values that correspond to specific*α*_*cr*_, *β*_*cr*_, *q*_*cr*_.

### 2.2 Non-null Bootstrapping for Single Inference

The above-mentioned procedure is applicable for large-scale multiple testing, as this enables the estimation of empirical mixed density *f* (*z*), the priors *p*_0_ and *p*_1_, and eventually the Posteriors *P*_0_(*z*) and *P*_1_(*z*). For single statistical testing (*N* = 1), similar stages can be followed in order to calculate the test statistic, estimate the histograms or probability density functions, and calculate the null distribution. However, it is not possible to estimate the priors; hence, these variables are not available. Notwithstanding, it is possible to estimate *f*_1_(*z*) by a different approach, namely the non-null bootstrapping which will make it possible to estimate *α* and *β* for a specific threshold, which would be a criterion on *z* (rather than on *P*_1_(*z*)). This approach is explained below:

#### 2.2.1 Non-Null Bootstrapping & Estimating *f*_1_ (*z*)

To estimate the non-null distribution *f*_1_(*z*) we may rely on bootstrapping when the grouping information of the data *g*_*i*_ are respected (in contrary to the commonly practised null bootstrapping that aims to find the null distribution where the grouping information/labels are not respected). For this purpose, the following re-sampling algorithm was used:

1. If *m*_*b*0_ is the number of observations in group *g* = 0, take *m*_*b*0_ samples (with substitution) from {*X*_i._|*g*_*i*_ = 0} to build Ξ_*b*0_.
2. If *m*_*b*1_ is the number of observations in group *g* = 1, (*m*_*b*0_ + *m*_*b*1_ = *m*), take *m*_*b*1_ samples (with substitution) from {*X*_i._|*g*_*i*_ = 1} to build Ξ_*b*1_.
3. find the *A* (AUC for ROC curve) and consequently the *z* value according to equation (4) using the obtained set of the bootstrapped data Ξ_*b*0_ and Ξ_*b*1_.
4. Repeat this procedure (1-3) for *B*_1_ times to obtain the needed samples of *z*_*b*_, *b* = 1…*B*_1_.

The GMM was then chosen to find the distribution of *z*_*b*_ values, which estimates *f*_1_(*z*).

#### 2.2.2 Estimating *α* and Power (1− *β*)

Similar to the calculation of *α* and *β* for a given *P*_*cr*_ value in (7) and (8), it is possible to use numerical integration to find the relationship between *α* and *β*. If *F*_0_(*z*) is the cumulative density function corresponding to *f*_0_(*z*), then for a given two-tail decision region *Z*(*α*),

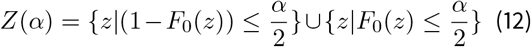

it is possible to describe *β* a as function of *α*:

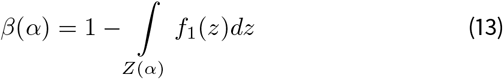

#### 2.2.3 Threshold Selection

The threshold selection in single testing is straight-forward, given the known relationship between *α* and *β* in (13). This can be pursued based on traditional *α* as thresholds, but now importantly based on specific values of *β* or power.

### 2.3 Numerical Implementation & Simulations

The numerical programming for the proposed EBI implementation was performed in MATLAB (versions 2016b-2018a, Mathworks Inc., Natick, MA, USA). The Empirical Bayesian Inference Toolbox for MATLAB is publicly available at https://github.com/NeuroMotor-org/EBI and is licensed under BSD 3-Clause ”New” or ”Revised” License .To demonstrate the utility of the proposed implementation and to test its validity, it was applied to simulated data. Simulated data allow comparison of the performance measures to real truth, which is not available in real life applications. All simulations were performed in MATLAB. Different simulations were carried out as detailed below.

#### 2.3.1 A Demonstrating Example

An example similar to applications in neural signal analysis and neuroimaging was considered. The simulation included *N*_*var*_ = 2000 variables, each with *m*_0_ = 20 observations/subjects in the first sample (e.g. controls), and *m*_1_ = 60 observations/subjects in second sample (e.g. patients), totalling *m_j_* = 80 observations/subjects. In control observations, all variables had a normal distribution 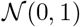, while in the other group the first 1600 variables had the same 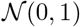 distribution, the next 300 variables came from 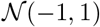 and the last 100 variables were from 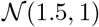.

#### 2.3.2 Comparison Against Real Truth

Using the simulated data from previous section, the *β* and *FDR* (or *q*) values were calculated as a function of the posterior threshold *P*_*cr*_ and were compared to the true values of *β* and *FDR*(*q*) at each *P*_*cr*_. The real values were found by using the original labels of the variables in the simulations. By comparing them to the detected labels by EBI, the true positive (TP), false positive (FP), false negative (FN) and true negative (TN) rates were calculated at each threshold, from which the real *β* and *FDR* were found.

Additionally, the same data underwent a similar analysis with a previous implementation of EBI (Efron, 2007b) in R (R Core Team, 2016), when used with the default value. Notice that this implementation of R, (which by default uses splines to estimate the density functions, and a parametric normal for the null distribution), is not guaranteed to converge for any data set in general (as it is fine-tuned for small prior *p*_1_ values) unless extensive manual tuning of parameters is used.

#### 2.3.3 Performance Under Different Condition

Several simulations were performed to test the performance of the framework in a broader range of conditions. This controlled variation of the simulation condition can test and inform of the performance in real-life applications, which is not easy to assess with typical experimental data due to the lack of a gold standard or real truth. The parameters for generation of simulated data and the application of the new EBI implementation included: *N*_*var*_ = 200, 2000, *p*_0_ = 0.25, 0.75, *m*_0_ = 25, 100, *m*_1_ = 25, 100, Normal (*σ* = 1) vs. Beta (*a* = 2,*b* = 10) distribution types, small or large difference/effect-size (Cohen’s *d* = 0.2, *d* = 0.9 for normal distributions; shift *d* = 0.05, *d* = 0.09 for Beta distributions). The estimated prior*p*_1_, real *FDR* at expected nominal value of 0.05, and real *β* at expected nominal value 0.2 were compared across all the 64 simulation conditions. Each simulation condition was repeated 3 times to account for the non-deterministic nature of the implemented bootstrapping and estimation procedures.

#### 2.3.4 Simulation of a Uni-Variate Example

To demonstrate the derivation of the *α* − *β* curve, a simple simulation with *m*_0_ = 15 and *m*_1_ = 25, random data with normal distributions for both groups (*σ* = 1), and shift value (Cohen’s *d*) of 0.5 was considered.

### 2.4 Exemplary Application on Experimental Data

The new implementation of EBI was applied on real-life experimental High-Density EEG data to further demonstrate the utility in practice. The experimental EEG was recorded during steady state conditions, including a resting-state condition (S. Dukic et al., 2017; Stefan Dukic et al., 2019) and a sustained isometric motor task (Coffey et al., 2019). The data included 363 and 99 epochs of 1 second duration in each condition respectively. Details of the spectral analysis have been previously reported (Nasseroleslami et al., 2019). EBI was used to find the significant difference between the rest and motor conditions over 128 EEG channels in 7 frequency bands that cover the range of 2-47Hz (7 *×* 128 = 896 variables).

## 3 Results

### 3.1 Example of Multiple Testing with EBI

Figure 1 exemplifies the generated results and a report for a typical simulated case as described in Section 2.3.1.Notice how the two probability density functions *f*_0_(*z*) and *f*_1_(*z*), as well as the prior *p*_1_ are the essential components in giving rise to the Posterior distributions *P*_0_(*z*) and *P*_1_(*z*). In addition pay attention to how the choice of different threshold levels (color-coded based on the criteria) helps to choose an informed statistical threshold for inference based on the levels of FDR and power they afford.

**Figure 1.**
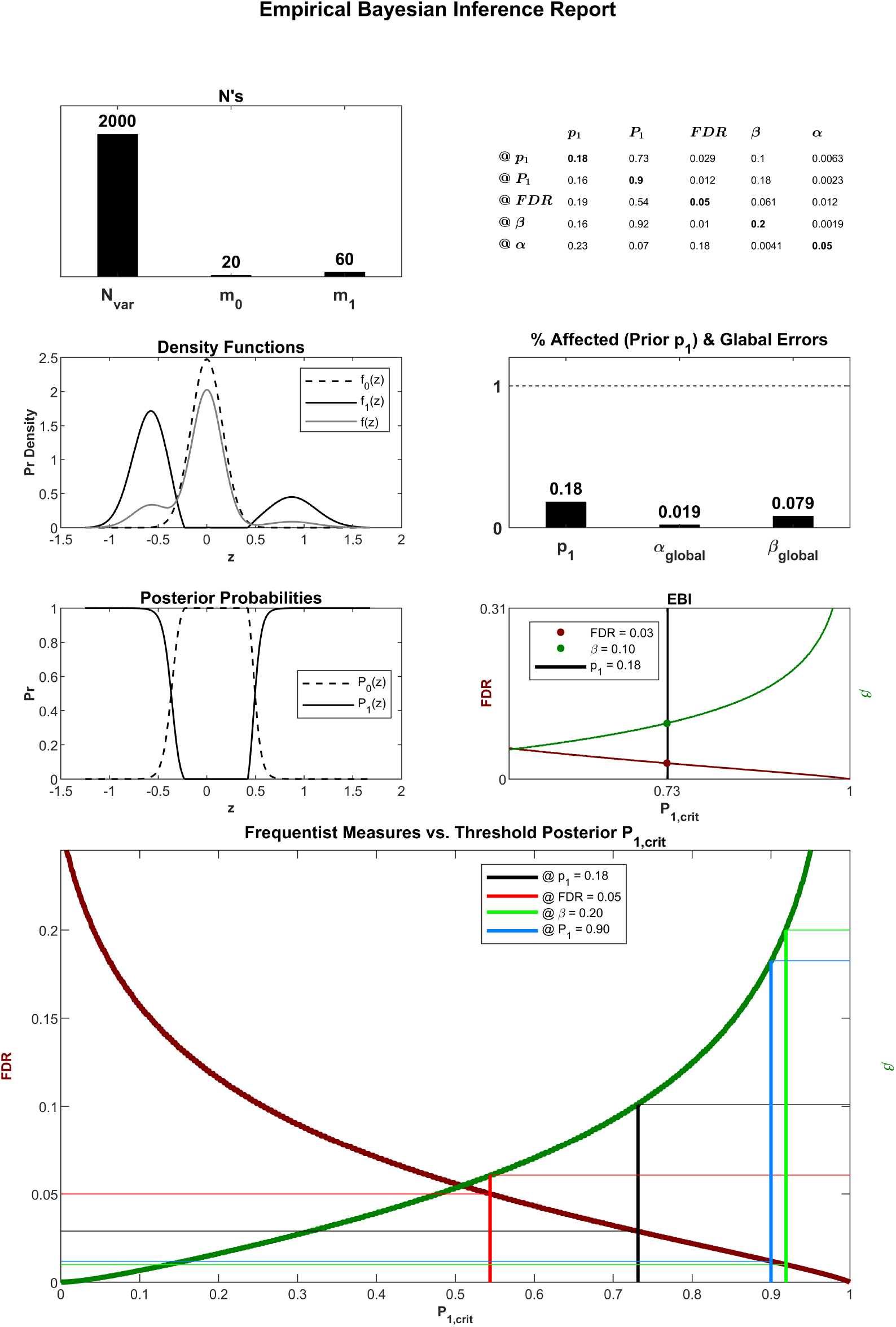
Report of an Empirical Bayesian Inference, applied on typical simulated data. The probability density functions of the mixed (*f* (*z*)) and null (*f*_0_(*z*)) data, estimated from the original and permuted, as well as of the non-null (*f*_1_(*z*)) data, estimated from *f* (*z*) and *f*_0_(*z*) by Bayesian inference are plotted as a function of the *z* (which is transformed to *z* from an original test statistic). Following the estimation of the fixed prior probability *p*_1_ = 0.19 (the ratio of affected to non-affected variables), the Posterior probabilities for null (*P*_0_(*z*)) and non-null (*P*_1_(*z*)) are estimated. The *α*_*global*_ = 0.02 and *β*_*global*_ = 0.083 show the global level of Type I and II errors regardless of a specific threshold, calculated from the probability density functions and Posterior Probabilities by numerical integration. The threshold selection is facilitated by the plots of FDR and *β* as a function of *P*_1,*crit*_. The table indicates common criteria as a function of *p*_1_, *P*_1_, *FDR*, *β* and *α* (thresholds are shown as bold diagonal values), and other corresponding values afforded by the selected criterion in each row. For example, at *p*_1_ = 0.18, the estimated *FDR* and *β* values will be 0.029 and 0.10 respectively. In the large plot, for each chosen criterion labelled by colour-coding in the legend, the projections to left and right axes indicate the afforded *FDR* and *β* by each criterion, respectively.

### 3.2 Comparison of Typical Behaviour Against Truth

Figure 2 compares the estimated FDR and *β* as a function of the threshold values (*P*_1_), against the real truth, using simulation labels as described in Section 2.3.2. Notice the similarity of real truth curves and estimates by the new implementation. In addition, as the simulated condition had a low prior *p*_1_, the results from the previous *locfdr* implementation in R could be calculated, which showed a good conformity to the real truth and the new implementation.

**Figure 2.**
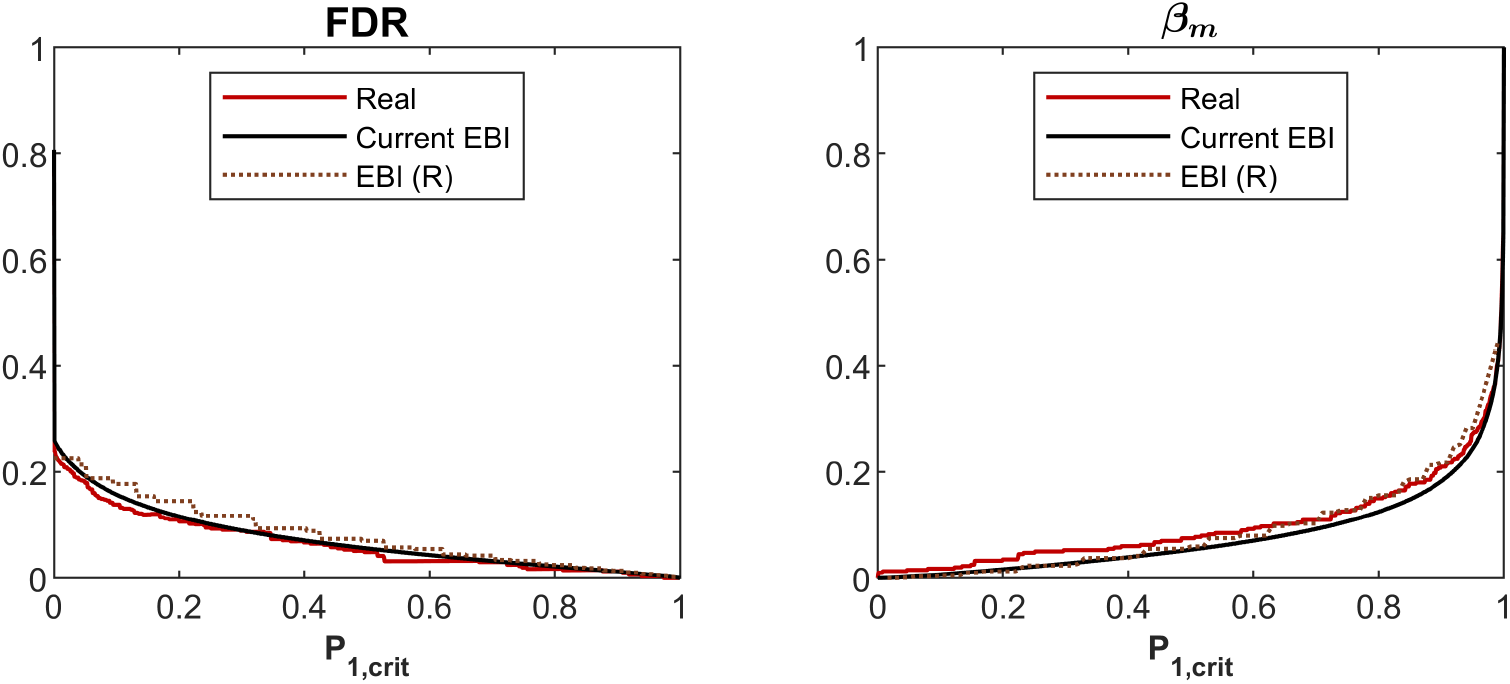
Comparison of FDR and *β* found by the EBI implemntation against the real truth in simulations. The data and conditions correspond to Figure 1. The dotted lines show the estimations from the *locfdr* package in R.

### 3.3 Performance of the EBI under Various Simulation Conditions

Figure 3 compares the estimated prior *p*_1_ against the real values, as well as the real FDR and *β* values at the selected threshold when estimated at nominal values at 0.05 and 0.2 (Described in Section 2.3.3). In the majority of conditions, the estimated measures were very close to the real values and there was negligible difference between the 5 different iterations of the simulation. The exception is at low effect sizes, combined with low number of observation/subjects and extreme prior values, where the estimation errors increase (possibly due to dissociations between the affected and non-affected variables and density functions). In the majority of the simulation cases the *locfdr* R package did not converge; therefore, the results were not included.

**Figure 3.**
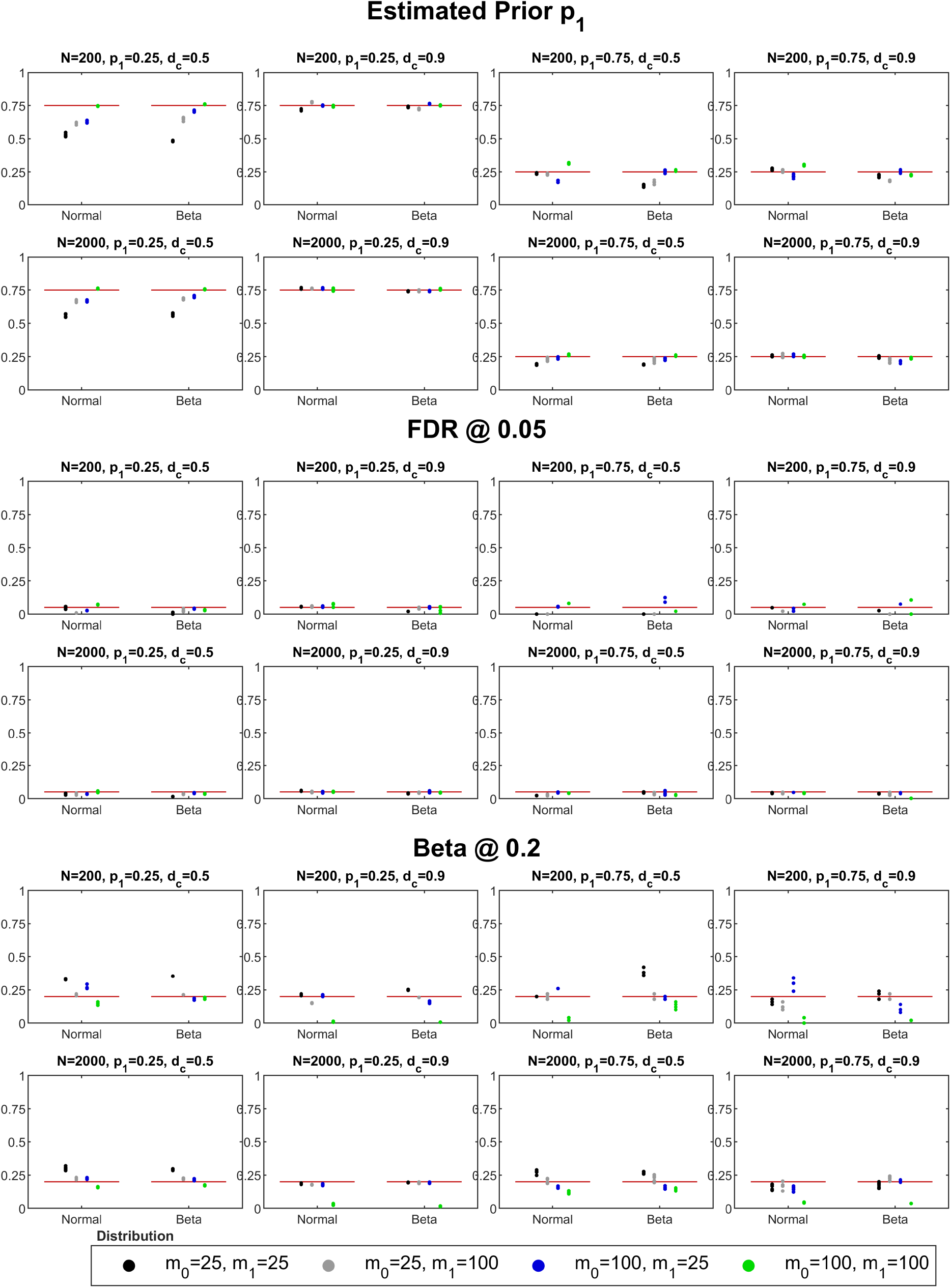
Comparison of estimated prior *p*_1_ against real truth, and observed FDR and *β* values at nominal values 0.05 and 0.2 of the selected thresholds in simulations. See Section 2.3.3 for details of methods. Real truth and nominal values are shown by red lines.

### 3.4 Example Single Testing

Figure 4 shows the correspondence of different *α* values to *β* values, for a an exemplary simulated data (Section 2.3.4).

**Figure 4.**
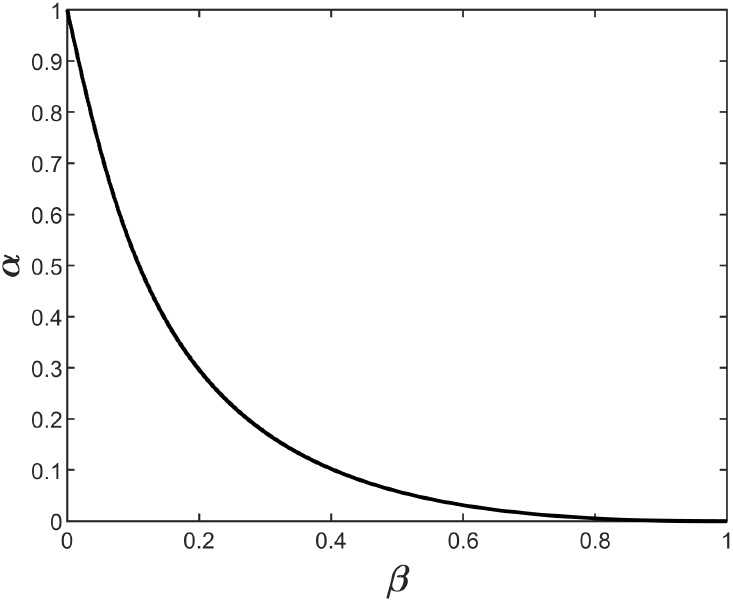
Correspondence of different *α* values to *β* values. The plot can be used for informed selection of threshold, similar to multivariate EBI.

**Figure 5.**
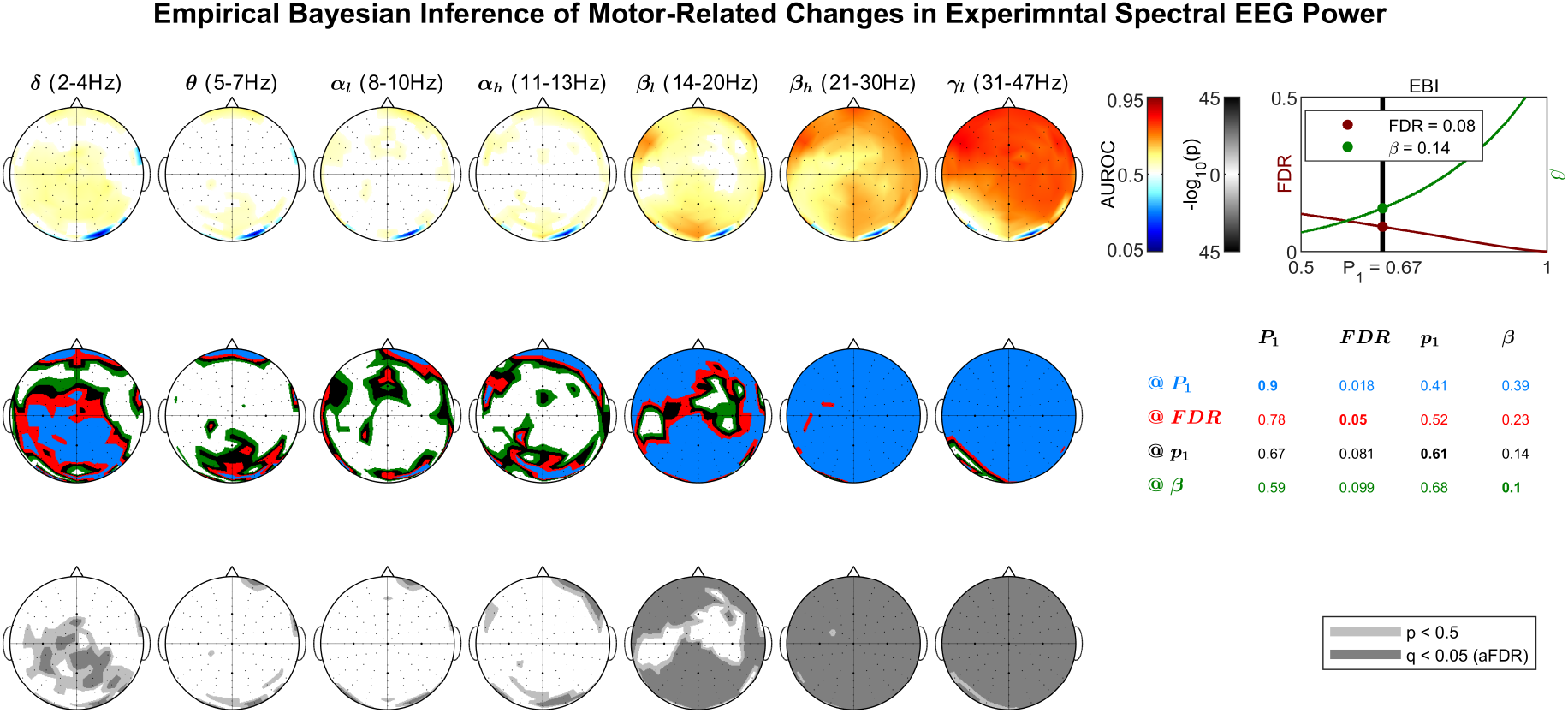
Threshold selection for the detection of significant change in spectral EEG power in steady-state, compared between rest and active motor task. The data corresponds to 1 subject and calculated from 363 and 99 epochs of 1 length during resting-state and isometric force generation, respectively..

### 3.5 Application on Experimental Data

Figure **??** shows the application of the EBI on experimental EEG data, showing the cortico-muscular coherence across EEG channels and 7 frequency bands. The effect of informed threshold selection on the experimental data and the corresponding values (Power, FDR, prior and Posterior) are demonstrated.

## 4 Discussion

While EBI (Efron, 2007b; Efron et al., 2001) provided a comprehensive theoretical framework for multivariate high-dimensional inference, the previous numerical implementation of EBI provided valid results only in limited conditions, namely, low prior *p*_1_ values and for rather high threshold values. Importantly, the previous implementation required numerous adjustments and parameter selection. The new implementation eliminates the need for parameter tuning (especially by using AIK for GMD fitting) and allows the method to be used in broader range of conditions. Importantly, the statistical power is explicitly estimated and made available for inference.

### 4.1 Applications

The new approach suits applications involving neural signal analysis, such as electromyography (EMG), Electroencephalography (EEG), as well as neuroimaging, e.g. Magnetic Resonance Imaging (MRI). More specifically, spectral, time-frequency, as well as functional and effective connectivity analyses can benefit most from the new statistical implementation. In applications such as fMRI, the need for improved statistical inference has been explicitly emphasised by highlighting the limitations with existing techniques that lead to high false discovery rates (Eklund, Nichols, & Knutsson, 2016). The existing attempts to improve the statistical inference in EEG connectivity analysis (Singh, Asoh, & Phillips, 2011; Singh et al., 2015), have yield only partial success to date. Here, we used simulations to compare the EBI reports against the real truth. Importantly, in 2 recent studies, we showed that EBI is reasonably cross-validated against traditional frequentist methods. EBI was cross-validated against the correction of significance level *α* according to the number of principal components in data (Iyer et al., 2017) when testing the significance of EEG time series. Moreover, the inference of EBI was cross-validated against adaptive False Discovery Rates (aFDR) (Benjamini & Hochberg, 1995; Benjamini et al., 2006) in comparing the average EEG connectivity patterns between healthy individuals and patient groups (Nasseroleslami et al., 2019).

A unique advantage of EBI is its ability to implicitly account for potential positive and negative correlations that may be present in the data. It is therefore a suitable candidate for situations where positive or negative correlations exist in multi-dimensional data (e.g. EEG/MEG network connectivity analysis or structural or functional MR imaging). This is afforded by the way the individual z-values pertaining to each variable are aggregated (i.e. the independent calculation of the test scores) and by the chosen approach for the calculation of a null distribution that similarly corresponds to the same data with or without correlation structures (Efron, 2007a). The flexible estimation of null distribution from permuted data by GMM affords such flexibility for inference in rather broad conditions.

In applications where only simple statistical testing is required, the calculated FDR is an accurate estimation equivalent to the procedures for pooled multivariate permutation tests, which can be used without reference to Bayesian inference.

#### 4.1.1 Practical Use

In practical application of single-variate statistical test, the p-values (The probability of data given the null hypothesis) are compared against a chosen threshold such as 0.05 or 0.001. While such established rules of thumb are not necessarily connected to the probabilities such as the Bayesian Poterior or AUC, they facilitate their practical uses in day-to-day application. The discussion of the appropriate thresholds for Bayesian Posterior, Power, and FDR are beyond this report. However, a brief discussion to make their meaning more accessible in the context of the applications shall make it easier to adopt rules of thumb for the thresholds.

- *P*_1_: For the Bayesian Posterior (The probability of effect given the data), any value larger than 0.5 should help to show the presence of an effect. However, in practice, values larger than 0.7, 0.8 and 0.9 can be better associated to small, medium and large effects.
- *FDR*: In multivariate applications, it is usually more informative to generalise the p-value using FDR, i.e. the probability/rate of false detection in the whole detection set, rather than by the *α*, i.e. the probability of having even a single false detection. For FDR or *q* (The ratio of the falsely detected variable to the all of the detected variables), the common sensible values should be interpreted based on the requirements of the applications. It is helpful to answer what percent of grime would be acceptable in figuring out the right pattern and extent of the effect in the variables. Usually the values of 0.25, 0.1 and 0.05 would be good first choices for a rough, adequate and good figure of the effects. Of course in situations where few false detection is affordable, lower values and the use of Type I error *α* as a better measure may be proffered. This however usually results in lower statistical power. It is useful to mention that while the *FDR* estimates the proportion of the falsely discovered effects, the experimenter/analyst can still put more weight and emphasis in the interpretations on the detections with higher Posterior *P*_1_ or AUC (as test statistic) values.
- Power (1−*β*): The multi-variate interpretation of the statistical power or 1 − *β* is the ratio of the correctly detected variables to all of the affected variables that exist. This can be similarly best interpreted in the context of applications. Finding a reasonable threshold can be facilitated by answering the question: What minimum percent of the full picture of the effect should the detection uncover? This is closely related to the reproducibility and replicability of the findings (single-variable interpretation of power). For example, if a study is powered at 0.5, only half of the really affected variables have been detected; therefore, in a future replication of the study, another 50% of the effects may be detected, leading to no apparent similarity if the results cannot be explained or aggregated by supplementary experiments or analyses. The statistical power of 0.7, 0.8, 0.9 can be very roughly considered as minimal, adequate and good levels in general.

Commonly a graphical representation of the FDR-Power curve (Fig. 1, bottom) is very helpful for an informed selection of the threshold based on the other performance measures afforded by each threshold. This actual meaning of these measures can greatly facilitate the conceptual interpretation of the thresholds and their actual utility in applications (see Figure 6 for a schematic ilustration). For example the table in Fig. 1 (top right) shows that when the threshold is chosen to in order to provide the same proportion of the affected/non-affected variables (to match the estimated prior probability, here 0.18), this would result in a Posterior Probability threshold of 0.73, a FDR of 0.029 and power of 1−0.1=0.9. It is also possible to define the threshold based on pre-determined criteria. For example choosing Posterior probability threshold of 0.9 results in detection of 16% of the variables (prior *p*_1_), FDR of 0.012, but a power of 1−0.18=0.82. Selection of FDR = 0.05 as threshold equals a posterior probability threshold of 0.54, which results in the detection of 19% of the variables and an estimated power of 1−0.061=0.939. Finally, if the threshold is chosen to afford a power of 1−0.2=0.8, this would correspond a Posterior probability threshold of 0.92, selection of 16% of variables, and FDR of 0.01. This provides the opportunity for informed selection of the threshold for specific applications.

**Figure 6.**
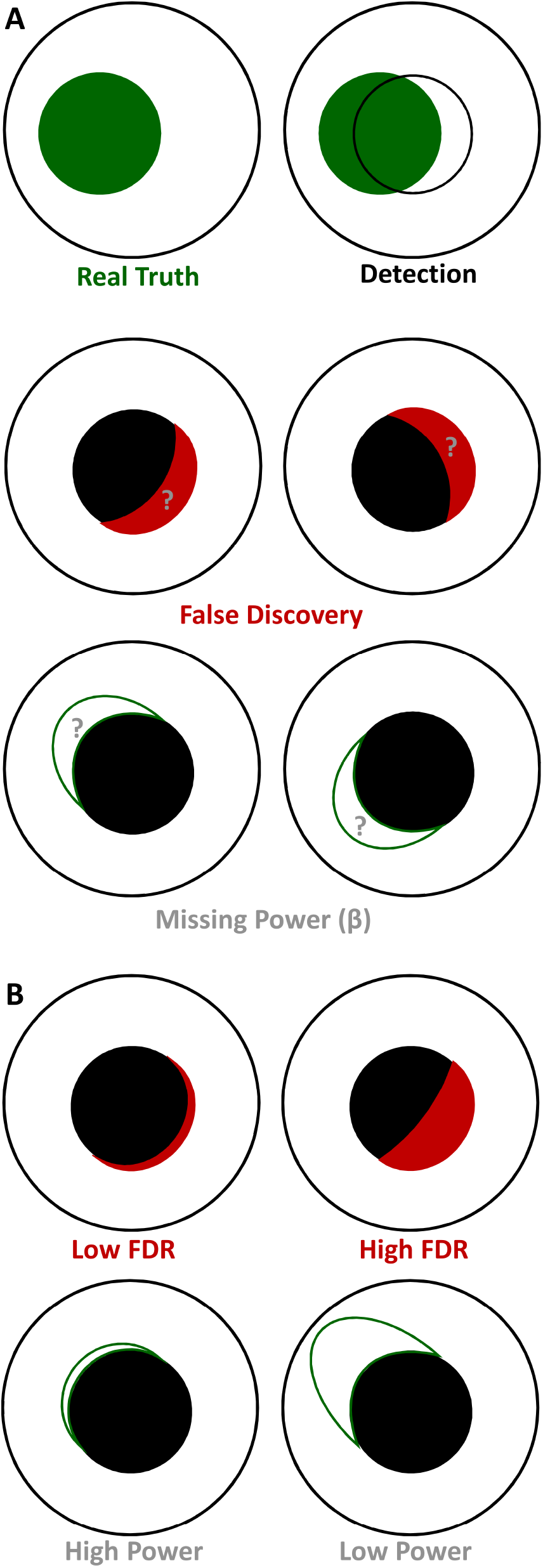
Schematic illustration of multivariate power and FDR in statistical inference. **A.** Illustration of the ground truth and the detections in statistical inferences such as EBI, where multivariate frequentist statistics can be estimated and used as thresholds. **B.** Demonstration of low vs. high values of FDR and Power, which can be chosen based on the requirements of scientific aplications and applied in EBI. Note that the size of the real truth corresponds to the (estimated) prior *p*_1_ and choosing a threshold at the same *p*_1_ means using a detection area as large as the real truth (see Figure 1).

### 4.2 Limitations

The practical range for the number of variables is between 100-10000. The performance beyond this range degrades. This limitation originates from the EBI framework, rather than a specific numerical implementation.

Too few variables lead to inaccurate probability density estimations where few isolated data points are not adequately represented by continuous distributions. In this situation, the extreme values of prior probabilities would correspond to fewer data points with real effect; hence, the probability densities fitted to these values will not be very representative and accurate.

On the other hand, too many variables lead to unwanted spread of the null distribution to the extent that inference at low FDR values does not yield significant results. This situation, however, can be partly remedied for by applying the EBI as several independent batches of analyses on the mutually exclusive chunks of data, each containing different variables. This is permissible as the quantities such as FDR and Posterior probability (and to a reasonable extent the power) are not affected by multiple testing (as is the case for p-values).

As the complete procedure for EBI relies on permutations for building the null distribution, the procedure would depend on random number generations and some variability in each run. Additionally, the numerical procedures for estimating the GMM fits to the distributions are subject to minor variability in each run. These 2 factors make the inference a non-deterministic procedure, which is subject to some variability. While important to take this into consideration, the results in Figure 3 idicate that this variability does not change the nature of the results.

Future studies are expected to focus on the factors that lead to inaccurate numerical estimations, further extending the range of operating conditions, as well as theoretical developments for robust estimation of prior when extreme data and conditions are processed.

### 4.3 Informed Selection of Appropriate Statistical Inference

Statistical inference from multi-dimensional data, especially the neural signals such as EEG/MEG, EMG, (fMRI) is challenging. It is crucial to take into consideration the potential advantages and disadvantages of each inference method, as well as the assumptions used in each inference framework to be able to choose the most appropriate method. The methods such as Statistical Parametric Mapping (SPM) (Friston et al., 1994) or Cluster-Based Permutation (CBP) (Bullmore et al., 1999; Maris et al., 2007) incorporate assumptions on the spatial distribution of the variables, therefore, the features showing the same effect within a spatial neighbourhood get an increase chance of detection. This is implemented by a smoothness assumption based on random field theory (Mehrkanoon, Breakspear, & Boonstra, 2014; Siegmund & Worsley, 1995; Worsley, 2001) in SPM, and by ”clustering” the variables based on a statistical threshold and added weights based on spatial adjacency in time, space and/or frequency, in CBP (Bullmore et al., 1999; Cohen, 2014; Maris et al., 2007). This assumption may help to better detect the effects in the groups of adjacent variables, but may not be optimal in detecting the boundaries of the effects or to detect all the components of the same effect with disjoint spatial presence. EBI does not impose any explicit spatial constraints in the inference from the variables. EBI implicitly accounts for such possible connections through the increased/accumulated probability of effect for such variables with similar test statistics that may originate from spatial adjacency or other correlations or interdependence in the data. This allows a better detection of the boundaries of the effect and equal chance of detecting the components of a spatially disjoint effect. However, the interpretation of the spatially isolated detection will be the job of the analyst or researcher.

## 5 Conclusion

The implementations of statistical inferences such as EBI that can inform of the posterior probabilities and statistical power need to be converted to common practice. This implementation of EBI and single testing which supports threshold selection has potential to add value to the neural signal analysis and neuroimaging studies by enabling realistic inference on high-dimensional multivariate data.

## Information Sharing Statement

The Empirical Bayesian Inference Toolbox for MATLAB is publicly available at https://github.com/NeuroMotor-org/EBI and is licensed under BSD 3-Clause ”New” or ”Revised” License. It includes the codes for statistical implementation, as well as the codes and processed data for the examples.

## Compliance with Ethical Standards

Conflict of Interest: The author declares that he has no conflict of interest.

## Acknowledgement

The author would like to thank the students and staff in the Academic Unit of Neurology, at Trinity College Dublin, the University of Dublin for facilitating and supporting this work. In particular, the useful comments on the manuscript by Stefan Dukic, Teresa Buxo, and Roisin Mc Mackin in the Academic Unit of Neurology and Muthuraman Muthuraman (Johannes Gutenberg-University, Mainz, Germany) are greatly appreciated. The study was supported by Irish Research Council (Government of Ireland Postdoctoral Research Fellowship GOIPD/2015/213 to the author), Science Foundation Ireland (SFI/16/ERCD/3854), and the Health Research Board of Ireland and the Irish Motor Neurone Disease Research Foundation (MRCG-2018-02).

## Appendix A: Extension of the Tests from Comparison to Correlation Coefficients and Beyond

The original EBI has been primarily used for two-sample one-dimensional location problems (between-group comparison) such as gene discovery by comparing a control group to a treatment or affected group, or similarly by comparison of the healthy individuals against patients as intended in neuro-electro-magnetic signal analysis. Notwithstanding, the framework can be similarly used for virtually any statistical test, such as one-sample location problems (where comparison of data against zero or paired comparison of data are intended), as well as correlation analysis. These options have been implemented in the EBI Toolbox for MATLAB.

## A1. One Sample Inference

For a one-sample 1-dimensional location problem, including *n* data points *x*_*i*_, the Wilcoxon’s Signed Rank test statistic is defined as 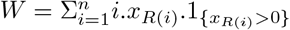, where *R*(*i*) is the rank of the {|*x*_*i*_|| *x*_*i*_ ≠ 0}. The function 1_{*x*}_ is 1 for the *x* that meet the condition, and 0 otherwise .The normalised test static may be defined as *W_n_* = *W* / (Σ*R*(*i*)) which is bounded between 0 and 1. Therefore, *W*_*n*_ can be transformed to the *z* space using (4), as in the case of AUC and the subsequent procedures will be similar to the two-sample problem.The bootstrapping procedure for building the null distribution, is carried out by performing random sign flips (multiplying data by random 1 or 1 values) and recalculation of the test statistics for the number of bootstrapping cycles.

## A2. Correlation Coefficient

To use the same framework for analysis of correlation coefficients, the Spearman’s correlation coefficient *ρ* can be mapped to the [0 1] range (as for AUROC, *A*) by the transformation (*ρ* + 1)/2. In this case, the grouping information will have equal number of paired zeros and ones. The null permutation of the data shall consist of using separate re-sampling (with substitution) from the first and second groups of observations for the same data points as the original data, which disregards their pairing information.

## References

Akaike, H. (1974, December). A new look at the statistical model identification. IEEE Transactions on Automatic Control, 19(6), 716–723. doi:10.1109/TAC.1974.1100705

Amrhein, V., Greenland, S., & McShane, B. (2019, March). Scientists rise up against statistical significance. Nature, 567(7748), 305. doi:10.1038/d41586-019-00857-9

Ashburner, J. (2012, August 15). SPM: A history. NeuroImage. 20 YEARS OF fMRI, 62(2), 791–800. doi:10.1016/j.neuroimage.2011.10.025

Benjamini, Y., & Hochberg, Y. (1995, January 1). Controlling the False Discovery Rate: A Practical and Powerful Approach to Multiple Testing. Journal of the Royal Statistical Society. Series B (Methodological), 57(1), 289–300.

Benjamini, Y., Krieger, A. M. & Yekutieli, D. (2006, January 9). Adaptive linear step-up procedures that control the false discovery rate. Biometrika, 93(3), 491–507. doi:10.1093/biomet/93.3.491

Bullmore, E. T., Suckling, J., Overmeyer, S., Rabe-Hesketh, S., Taylor, E., & Brammer, M. J. (1999, January). Global, voxel, and cluster tests, by theory and permutation, for a difference between two groups of structural MR images of the brain. IEEE Transactions on Medical Imaging, 18(1), 32–42. doi:10.1109/42.750253

Coffey, A., Buxo, T., Fasano, A., Dukic, S., Heverin, M., Lowery, M., … Hardiman, O. (2019). Cortico-Muscular Coherence Patterns in Motor Neuron Disease. (p. 5). European Network for the Cure of ALS 2019 Meeting. Tours, France.

Cohen, M. X. (2014, January 17). Analyzing Neural Time Series Data: Theory and Practice (1 edition). Cambridge, Massachusetts: The MIT Press.

Dukic, S. [S]., Iyer, P. M., Mohr, K., Hardiman, O., Lalor, E. C., & Nasseroleslami, B. (2017, July). Estimation of coherence using the median is robust against EEG artefacts. In 2017 39th Annual International Conference of the IEEE Engineering in Medicine and Biology Society (EMBC) (pp. 3949–3952). 2017 39th Annual International Conference of the IEEE Engineering in Medicine and Biology Society (EMBC). doi:10.1109/EMBC.2017.8037720

Dukic, S. [Stefan], McMackin, R., Buxo, T., Fasano, A., Chipika, R., Pinto-Grau, M., … Nasseroleslami, B. (2019, July 26). Patterned functional network disruption in amyotrophic lateral sclerosis. Human Brain Mapping, 0(0), 1–16. doi:10.1002/hbm.24740

Efron, B. (2004). Large-Scale Simultaneous Hypothesis Testing: The Choice of a Null Hypothesis. Journal of the American Statistical Association, 99(465), 96–104.

Efron, B. (2007a). Correlation and Large-Scale Simultaneous Significance Testing. Journal of the American Statistical Association, 102(477), 93–103.

Efron, B. (2007b, August). Size, power and false discovery rates. The Annals of Statistics, 35(4), 1351–1377. doi:10.1214/009053606000001460

Efron, B., Tibshirani, R., Storey, J. D., & Tusher, V. (2001). Empirical Bayes analysis of a microarray experiment. Journal of the American statistical association, 96(456), 1151–1160. doi:10.1198/016214501753382129

Eklund, A., Nichols, T. E., & Knutsson, H. (2016, December 7). Cluster failure: Why fMRI inferences for spatial extent have inflated false-positive rates. Proceedings of the National Academy of Sciences, 113(28), 7900–7905. doi:10.1073/pnas.1602413113. pmid: 27357684

Fisher, R. A. (1915). Frequency Distribution of the Values of the Correlation Coefficient in Samples from an Indefinitely Large Population. Biometrika, 10(4), 507–521. doi:10.2307/2331838.JSTOR:2331838

Friston, K. J., Holmes, A. P., Worsley, K. J., Poline, J.-P., Frith, C. D., & Frackowiak, R. S. J. (1994). Statistical parametric maps in functional imaging: A general linear approach. Human Brain Mapping, 2(4), 189–210. doi:10.1002/hbm.460020402

Iyer, P. M., Mohr, K., Broderick, M., Gavin, B., Burke, T., Bede, P., … Vajda, A. (2017). Mismatch negativity as an indicator of cognitive sub-Domain Dysfunction in amyotrophic lateral sclerosis. Frontiers in Neurology, 8, 395.

Le, C. T., Pan, W., & Lin, J. (2003, July 1). A mixture model approach to detecting differentially expressed genes with microarray data. Functional & Integrative Genomics, 3(3), 117–124. doi:10.1007/s10142-003-0085-7

Maris, E., Schoffelen, J.-M., & Fries, P. (2007, June 15). Nonparametric statistical testing of coherence differences. Journal of Neuroscience Methods, 163(1), 161–175. doi:10.1016/j.jneumeth.2007.02.011

McLachlan, G., & Peel, D. (2004, April 5). Finite Mixture Models. John Wiley & Sons.

Mehrkanoon, S., Breakspear, M., & Boonstra, T. W. (2014, October 15). The reorganization of corticomuscular coherence during a transition between sensorimotor states. NeuroImage, 100, 692–702. doi:10.1016/j.neuroimage.2014.06.050

Moore, D., McCabe, G., & Craig, B. (2010). Introduction to the Practice of Statistics. Macmillan Higher Education.

Nasseroleslami, B., Dukic, S., Broderick, M., Mohr, K., Schuster, C., Gavin, B., … Hardiman, O. (2019, January 1). Characteristic Increases in EEG Connectivity Correlate With Changes of Structural MRI in Amyotrophic Lateral Sclerosis. Cerebral Cortex, 29(1), 27–41. doi:10.1093/cercor/bhx301

Nuzzo, R. (2014). Statistical errors. Nature, 506(7487), 150–152.

Oostenveld, R., Fries, P., Maris, E., & Schoffelen, J.-M. (2010, December 23). FieldTrip: Open Source Software for Advanced Analysis of MEG, EEG, and Invasive Electrophysiological Data. Computational Intelligence and Neuroscience, 2011, e156869. doi:10.1155/2011/156869. pmid: 21253357

Qin, G., & Hotilovac, L. (2007, August 14). Comparison of non-parametric confidence intervals for the area under the ROC curve of a continuous-scale diagnostic test. Statistical Methods in Medical Research, 17(2), 207–221. doi:10.1177/0962280207087173

R Core Team. (2016). R: A Language and Environment for Statistical Computing. Vienna, Austria: R Foundation for Statistical Computing. Retrieved from https://www.R-project.org

Siegmund, D. O., & Worsley, K. J. (1995, April). Testing for a Signal with Unknown Location and Scale in a Stationary Gaussian Random Field. The Annals of Statistics, 23(2), 608–639. doi:10.1214/aos/1176324539

Singh, A. K., Asoh, H., & Phillips, S. (2011, September 1). Optimal detection of functional connectivity from high-dimensional EEG synchrony data. NeuroImage, 58(1), 148–156. doi:10.1016/j.neuroimage.2011.05.082

Singh, A. K., Asoh, H., Takeda, Y., & Phillips, S. (2015, March 30). Statistical Detection of EEG Synchrony Using Empirical Bayesian Inference. PLoS ONE, 10(3), e0121795. doi:10.1371/journal.pone.0121795

Worsley, K. J. (2001, December). Testing for signals with unknown location and scale in a χ2 random field, with an application to fMRI. Advancesin Applied Probability, 33(4), 773–793. doi:10.1239/aap/1011994029

Zhou, X.-H., McClish, D. K., & Obuchowski, N. A. (2009, September 25). Statistical Methods in Diagnostic Medicine. Hoboken, NJ, USA: John Wiley & Sons.

